# Chip-seq and gene expression data for the identification of functional sub-pathways: a proof of concept in lung cancer

**DOI:** 10.1101/2020.06.15.151712

**Authors:** Xanthoula Atsalaki, Lefteris Koumakis, George Potamias, Manolis Tsiknakis

## Abstract

High-throughput technologies, such as chromatin immunoprecipitation (ChIP) with massively parallel sequencing (ChIP-seq) have enabled cost and time efficient generation of immense amount of genome data. The advent of advanced sequencing techniques allowed biologists and bioinformaticians to investigate biological aspects of cell function and understand or reveal unexplored disease etiologies. Systems biology attempts to formulate the molecular mechanisms in mathematical models and one of the most important areas is the gene regulatory networks (GRNs), a collection of DNA segments that somehow interact with each other. GRNs incorporate valuable information about molecular targets that can be corellated to specific phenotype.

In our study we highlight the need to develop new explorative tools and approaches for the integration of different types of -omics data such as ChIP-seq and GRNs using pathway analysis methodologies. We present an integrative approach for ChIP-seq and gene expression data on GRNs. Using public microarray expression samples for lung cancer and healthy subjects along with the KEGG human gene regulatory networks, we identified ways to disrupt functional sub-pathways on lung cancer with the aid of CTCF ChIP-seq data, as a proof of concept.We expect that such a systems biology pipeline could assist researchers to identify corellations and causality of transcription factors over functional or disrupted biological sub-pathways.

## 1. Introduction

Gene regulatory networks are maps of genes that provide, among others, information of their interactions. Transcription factors are proteins that regulate the transcription of their target genes to produce messenger RNA, while in post-transcriptional regulation microRNAs (miRNAs) cause degradation and repression of target mRNAs. All these regulations can enrich a GRNs with link/edges from TF or miRNA genes to their target mRNAs. All these interactions are static, so we can represent a GRN with a map even though regulatory interactions occur dynamically in space and time [1]. Interactions in a GRN could be of many types such as activation, inhibition, catalysis, binds to and co-cited. Analysis of GRNs provides insight on the interactions between proteins/genes and the quantitative modeling of expression data can assist the identification of novel molecular approaches to target biological processes [2] or infer/provide new knowledge [3]. Many studies in the literature that deal with expression data and GRNs concern the reconstruction of networks using expression data [4]. Apart from the GRN reconstruction, many methodologies take advantage of the GRNs to group genes or select genes [1], [5]. The most advanced methods that combine GRNs and expression data take advantage of the semantics in a GRN such as the interaction of the genes and identify core regulatory genes that are potential targets for therapeutic intervention [6], [7]. GRNs have been well studied in the bacteria and yeast [8] and also in more complex organism such as humans even if it is challenging for a variety of reasons. Starting from the network per se, the gene to gene regulatory interactions are far from complete [9]. Furthermore, it is known that interactions vary across different types of tissues[8]. Another limitation of GRNs for higher organisms, is complexity since each gene/molecule has many targets and is also targeted by multiple regulators. There are also cases of self-regulation (auto-activation or auto-inhibition) or cross-regulation among molecules. Such difficulties make the human GRNs challenging not only to be produced but also to be analyzed than a simpler network such as the yeast GRNs [10].

The evolution of high-throughput and Next Generation Sequencing geared researchers to study biological systems at an elevated level [11]. Part of this evolution is the ChIP-seq that is the standard for the research of Protein-DNA interactions. ChIP-seq is a next generation sequencing technology for chromatin immunoprecipitation that provides valuable information for the transcription factors and their effect on phenotype specific mechanisms. Chip-seq is considered as a powerful mechanism to identify binding sites of DNA-associated proteins, such as Transcription Factors (TFs) and provides insights for a particular protein in living cells. The process includes the enrichment of specific DNA-protein complexes with the aid of an antibody. CHIP-seq experiments give us the ability to better understand the transcription factor biology, chromatin modification and transcription. Transcription factors and epigenetic modifications provide the basis for profiling transcriptional or epigenetic regulatory relationships, while binding features can be used to identify regulatory modules[12].

In the literature we can find reports that TFs take part in many important biological functions and human diseases, such as cell differentiation, proliferation, immune response, apoptosis, cardiac diseases, and tumor development [13]. Interpreting the regulation of TFs is helpful for understanding their regulatory function in complex biological systems. Moreover, biological pathways that represent complex interactions between proteins in living cells (GRNs) can help biologists to infer new knowledge for the biological interactions in a molecular level. Coupling GRNs and gene expression data can reveal molecular regulations in treatment response [14], [15]. Combining NGS (ChIP-seq) and GRNs can help the biologists to discover gene pathways associated with expression of genes in different phenotypes and help them understand complex biological processes such as cell cycle, cell differentiation, cell apoptosis, diseases. As a result, through the combination of a pathway analysis and the TFs that may disrupt key pathways for the specific disease, researchers can discover new treatments [16] while ChIP-seq can provide information related to the regulation of specific genes [17].

Even though ChIP-seq is a powerful sequencing methodology that has greatly advanced our understanding of chromatin and enhancer structures, in the literature there is limited research that combine regulatory networks and ChIP-seq. Recently, the Signaling Pathways Project [18] initiative provided a database that integrates transcriptomic datasets with biocurated ChIP-Seq datasets. The system provides also a meta-analysis technique that searches transcriptomic datasets and is able to provide ranked signatures, which allow for prediction of signaling pathway node-target regulatory relationships. ChIPBase [19] is also a database that combines non-coding RNAs and protein-coding genes aiming to identify transcriptional regulatory relationships between transcription factors and genes. Mars et al [20] proposed a numerical deconvolution approach that has been proved to improve both the resolution and quantitation of protein– DNA interaction maps from ChIP-Seq data. Xu et al [21] using ChIP-seq and expression data identified that the Bromodomain-containing 7 (BRD7) tumor suppressor gene contributes to the enrichment of the cell cycle and apoptosis pathway genes. Authors also constructed a regulating network of BRD7 downstream genes that revealed multiple feedback regulations. Malhotra et al [22] integrated ChIP-Seq with expression data and identified 645 basal and 654 inducible direct targets of Nrf2 while results were confirmed in an in vivo stress model of cigarette smoke-exposed mice. Martin et al [23] identified new genes involved in bZIP10 and conferred oxidative stress tolerance using ChIP-Seq from BdbZIP10. Raj et al [24] identified dysregulated key genes in breast cancer using ChIP-seq data and combined them with known protein-protein interaction networks for the selection of the most promising genes. In ovarian cancer, Kumar et al [25] used ChIP-Seq data along with protein-protein interaction network analysis and identified that PAX2, PAX5, FOXP1 and KLF16 are promising genes whose presence among differential peaks led to the positive conclusion of their role in the specific disease.

In this study, we assess the predictive power of ChIP-seq in conjunction with a differential sub-pathway analysis methodology that takes into account the regulations of genes and the binding sites of ChIp-seq data. We combined ChIP-seq data from the transcription factor CTCF on lung cancer cell line A549, derived from a Shiny application implemented for the purpose of this study and expression results from the MinePath [26] pathway analysis tool. Our analysis in lung cell cancer, identified a disrupted sub-path in MAPK pathway that in turn disrupts p53 signaling pathway; an indication that supports the vision of integrating different omics data can enhance the biological knowledge and disease treatment.

## 2. METHODS

Lung cancer is the leading cause of deaths from cancer. The resistance to therapy (radiotherapy or chemotherapy) is one of the issues that researchers try to address in the treatment of lung cancer patients. Contrary to other cancer types, biomarkers of lung cancer are still not applicable in the clinical practice guidelines. Most of the radio and chemo therapies use agents for lung cancer that target directly DNA. Since p53 functions as a hub gene for many biological processes including DNA repair, cell cycle and apoptosis, constitutes it as a perfect candidate for targeted therapy. p53 can be either upregulated/activated forcing cancer cells to be killed or downregulated/deactivated normal cells for temporary chemoradiation protection [14]. CTCF is a 11-zinc finger protein involved in many cellular processes such as transcriptional regulation, alternative splicing, regulation of chromatin architecture insulation, imprinting [27]. CTCF also regulates indirectly the p53 gene since there is aninteraction with the Wrap53, the natural antisense transcript of p53 and one of the target sites for the tumor suppressor protein p53 [28].

We conducted an experiment with the transcriptional repressor CTCF in lung cancer using pathway analysis from MinePath along with ChIP-seq-data from the Shiny application. Specifically, we used the Shiny application to infer the binding sites of CTCF on A549 cell line. We inserted the binding sites they were derived in MinePath. For the feasibility study we chose a dataset of 60 lung cancer samples and 60 healthy samples (GSE19804 from Gene Expression Omnibus). The transcriptome analysis of the gene expression data (60 lung cancer samples versus 60 healthy samples) conducted with MinePath taking into consideration all the human KEGG[29] pathways (299 in total).

### 2.1. ChIP-seq

We implemented a Shiny application that downloads ChIP-seq data from the ENCODE ChIP-seq Experiment Matrix [30]. We created a web interface (shinny application) in the statistical programming language R where the user (i) selects, (ii) analyzes the equivalent ChIP-seq peak files obtained from ENCODE Project and (iii) gets a file with the most significant binding sites. The file reports: (i) the gene id’s (EntrezID), (ii) the p-value, (iii) the q-value (or False Discovery Rate, FDR), (iv) the signal value and (v) the gene name. EntrezID has been selected as it is a genes coding system used also from the KEGG database. Signal value is the measurement of overall (usually average) enrichment for the region, which is the average (between replicates) read count over the region. If the signal value is high, it means that a lot of chromatin from that region is pulled down by the IP and sequenced.

The application chooses the optimal IDR (Irreproducible Discovery Rate) threshold peak files (IDR<0.01) and identifies all the binding sites near Transcription Start Sites (TSS) and within 5kb upstream of TSS within the gene. Transcription factors are proteins that bind to DNA, typically upstream from and close to the transcription start site of a gene, and regulate the expression of that gene by activating or inhibiting the transcription machinery [19]. We threshold the q-value of the peaks and we choose an FDR cut-off<0.01, since 1% FDR is the most commonly accepted value for peaks of good quality [12]. We map the genes with their corresponded Entrez Id’s using the Bioconductor Mapping Library EnsDb.Hsapiens and we get the selected TF’s binding sites for the specific cell line.

An overview of the pipeline is shown in Figure 1. We have also added one more functionality in the Shiny application that allows the user to intersect two different ChIP-seq file peaks and receive a file with all the overlapping peaks between these files for further differential analysis study.

**Figure 1.**
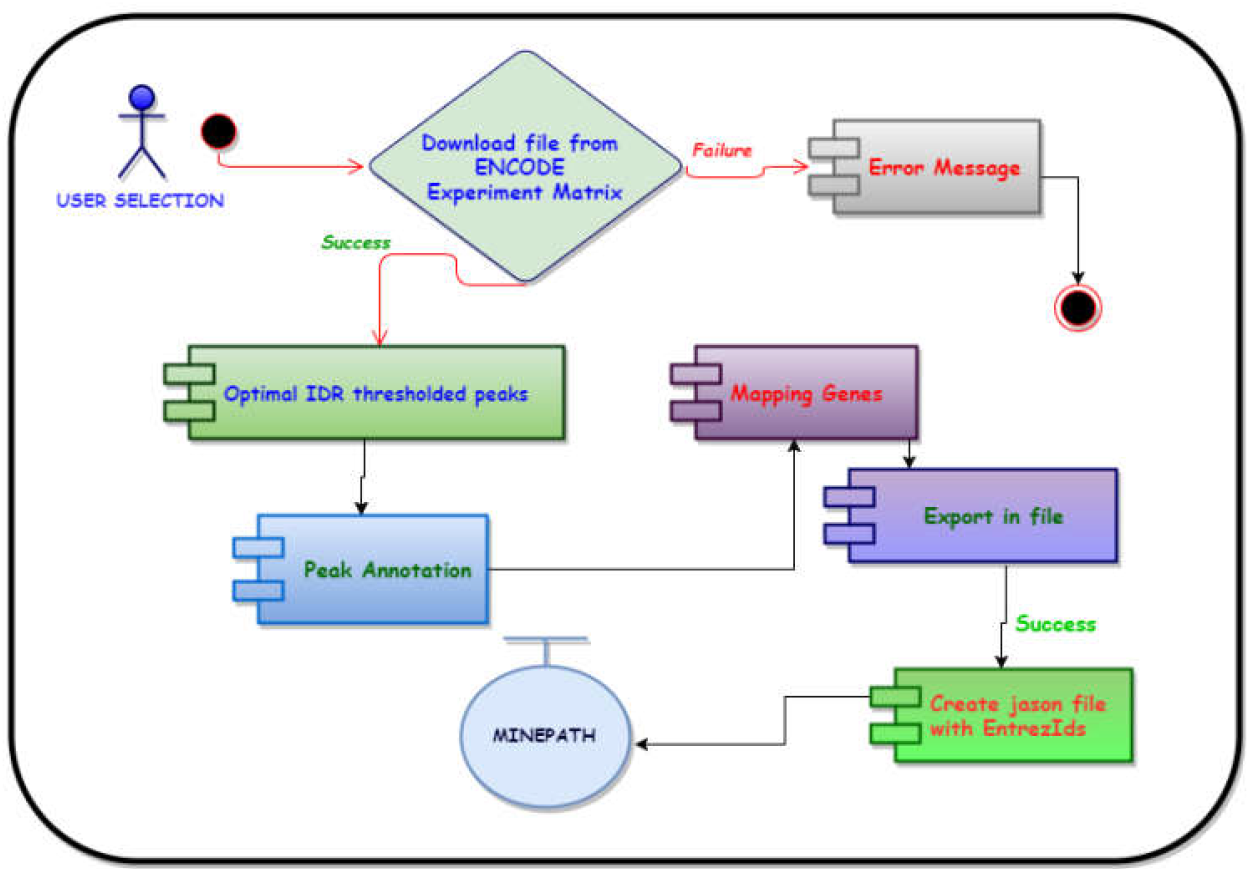
Experiment pipeline

### 2.2. Pathway analysis

MinePath is a pathway analysis tool that extract information from GRNs and identifies differentially expressed sub-pathways using gene expression data (e.g. mRNA, RNA-seq). For our methodology we also used and extended the open source GRN visualization tool, part of the MinePath pathway analysis web server (www.minepath.org), to accept and visualize ChIP-seq peak data files inferred from the Shiny application. For our experiment we used the binding sites of CTCF in A549 lung cancer cell line that we derived from our Shiny application while the algorithmic process of the identification of functional subpathways comes from MinePath.

## 3. RESULTS

According to MinePath the most significant pathway for the differentially expressed sub-pathways based on the gene expression data is the p53 with a p-value less than 10-8. Looking at the p53 pathway (Figure 2) in the MinePath viewer [31] we can conclude that p53 pathway is mainly functional for lung cancer samples (green relation between genes) as we expected.

**Figure 2.**
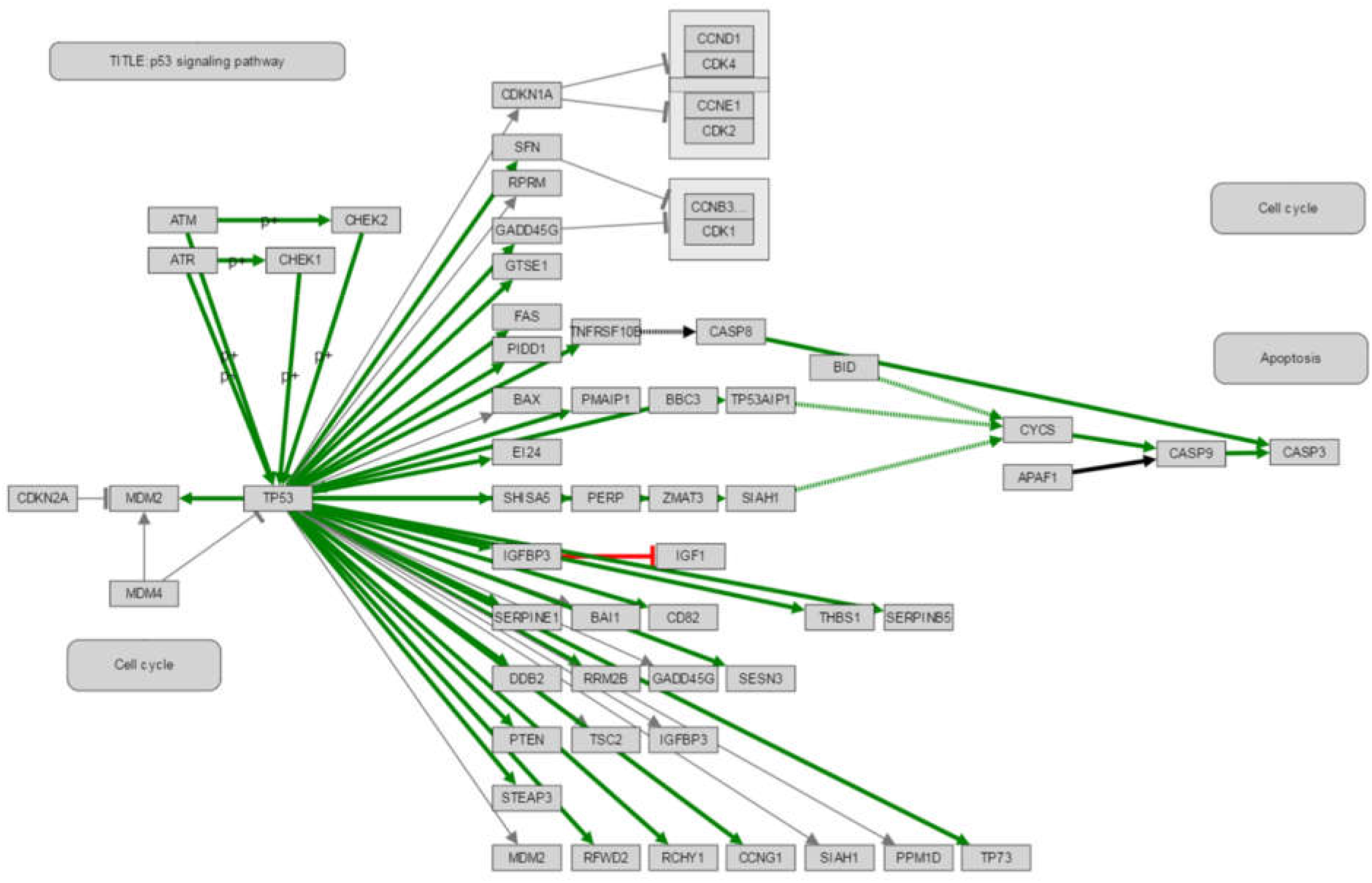
p53 signaling pathway

A research question would be “***What alerted p53 pathway only for lung cancer samples?***” One of the known cancer related roots, which can alter the p53 pathway, can be found in the mitogen-activated protein kinase (MAPK) signaling pathway. TP53 gene that plays a key role in p53 pathway is expressed in MAPK pathway. The MAPK pathway cascade is a highly conserved module that is involved in various cellular functions, including cell proliferation, differentiation and migration. MAPK regulates many transcription factors such as CREB, c-myc, C-Fos, while defects in the regulation of MAPK are known to contribute to uncontrolled growth in cells and cancer. MAPK, as well as other signaling pathways, can be disrupted if its genes are mutated and these genes can become stuck in the “on” or “off” position. Drugs that reverse the “on” or “off” switch are being investigated as treatments in various diseases including cancer [32].

As a result, we have chosen MAPK signaling pathway, one of the top ranked differential pathways with p-value <0.05 based on the results of MinePath. In Figure 3 we can view the functional and non-functional sub-pathways for lung cancer and healthy samples, red and green colors in edges respectively, along with the binding sites of the CTCF, red color on nodes, for the MAPK signaling pathway.

**Figure 3.**
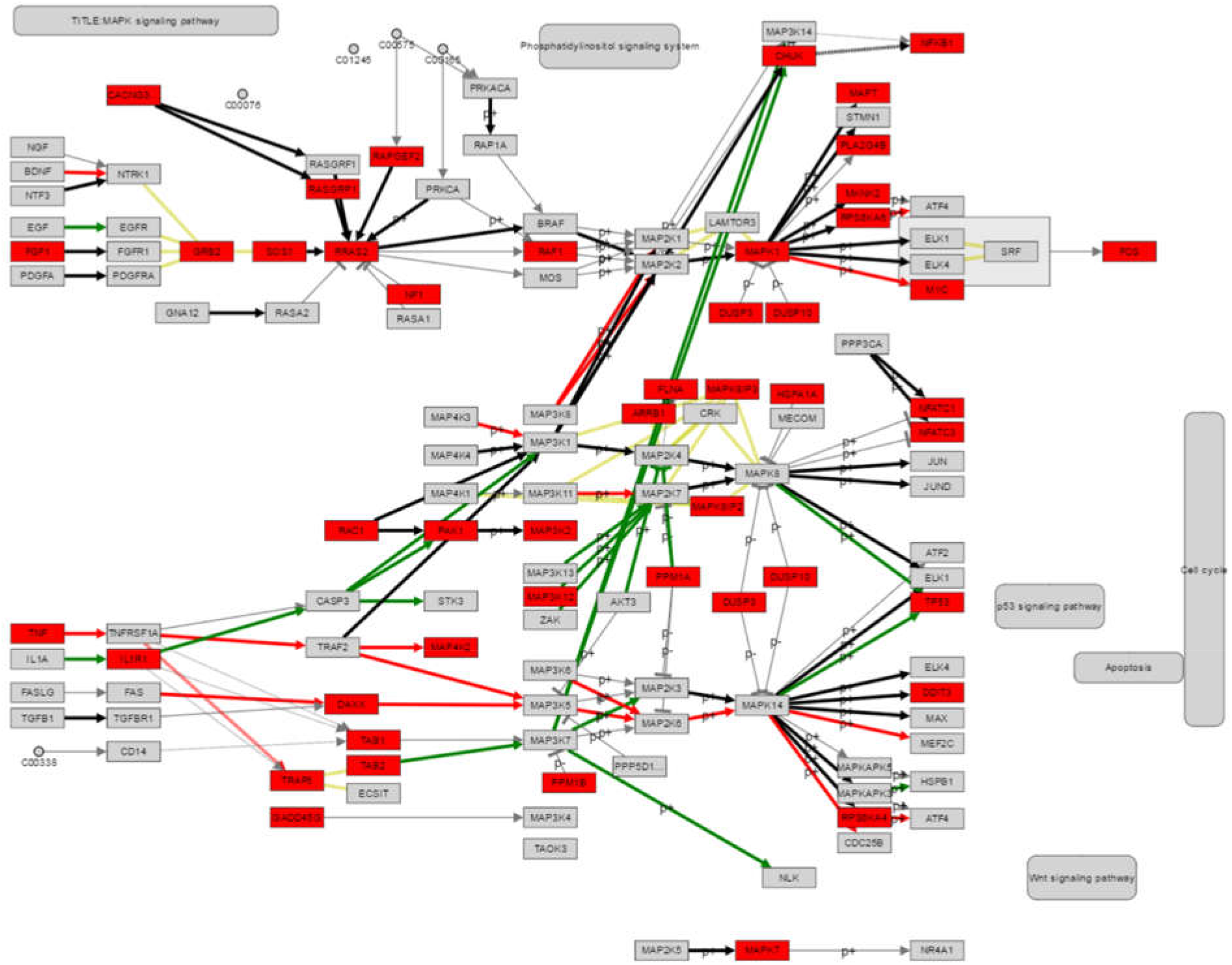
MAPK functional sub-pathways and binding sites of CTCF on A549

As we can see in Figure 2 Figure 3, GRNs are represented as networks where the nodes are genes and the edges relations between genes. The red nodes in our visualization show the genes that are affected from the transcription factor CTCF, based on our ChIP-seq pipeline analysis. The color coding for the edges follows the MinePath principles where green lines are the functional relations/reaction between genes, identified after the analysis of the expression data, for the first phenotype (in our case the lung cancer samplesļ), red are the functional relations for the second phenotype (healthy samples), grey the inactive relations and black the relations that are functional for both phenotypes (lung cancer and healthy samples). Finally yellow represent association/disassociation relations, which are considered physical associations and always hold Figure 3 highlights a part of the MAPK GRN, which shows a functional sub-pathway that starts from TAB2, activates MAP3K7, which alters MAP2K3 then MAPK14, and in turn activates TP53, which continue to the alteration of p53 signaling pathway, apoptosis and alteration of the cell cycle. Given the high p53 mutation frequency in lung cancer which likely impairs some of the p53-mediated functions, a role of p53 as a predictive marker for treatment responses has been suggested [33].

**Figure 4.**
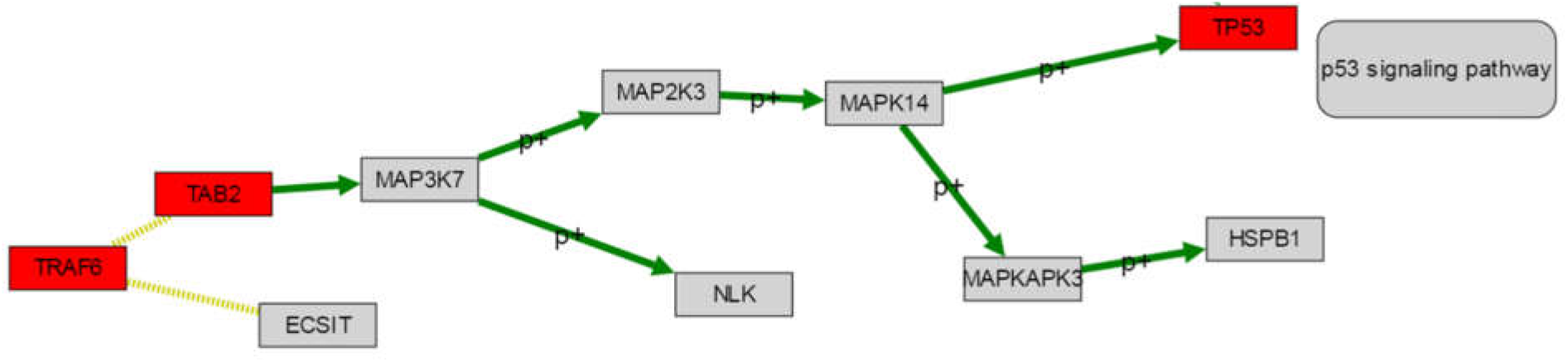
TAB2-TP53 Sub-path for lung cancer samples

## 4. DISCUSSION

Having in our disposal the ChIP-seq CTCF in lung cancer cell line data we can identify it’s binding sites and with the technology of pathway analysis we can target functional sub-pathways which lead to known oncogenic processes. Specifically we identified that we can disrupt a functional sub-pathway targeting the tp53 gene in MAPK, a pathway common among cancers that also play a key role in cancer treatment [32], since the binding sites of CTCF affect the TAB2, a trigger for a functional sub-pathway to the activation of p53. As a result, disruption of tp53 could lead to disruption of p53 signaling pathway that in turn affects apoptosis and cell cycle [28]. Only recently the TAB2 protein was associated with cancer disease and specifically as in a study that used Tamoxifen as endocrine therapy in breast [34]. Of course such evidence (from bioinformatics analysis and literature) needs to be biologically validated in vitro before we can safely associate the TAB2 protein with lung cancer. Furthermore, we have to acknowledge that our analysis is focused only in the affected pathways that MinePath identified but biological networks provide much more information that can follow up research outcomes such as the identification or reconstruction of a protein– protein interaction network [35], [36]. We also plan to provide the pipeline in the OpenBio-C workflow environment [37] as a bioinformatics pipeline proof of concept for the validity of integrating ChIP-seq and RNA-seq datasets.

Concluding, we expect that the integration of ChIP-seq, gene expression and gene regularity networks, can provide valuable hidden information for the research community and reveal links between transcription factors and biological sub-pathways that are known targets for disease treatments. We believe that the proposed automated methodology could provide insights in other domains also such as the systems pharmacology and empower the clinical decision support for personalized oncology [38].

## Acknowledgments

The work presented in this paper is supported by the OpenBio-C project (www.openbio.eu), which is cofinanced by the European Union and Greek national funds through the Operational Program Competitiveness, Entrepreneurship and Innovation, under the call RESEARCH – CREATE – INNOVATE (project id: T1EDK-05275).

